# Redistribution of FLAgellar Member 8 during the trypanosome life cycle: consequences for cell fate prediction

**DOI:** 10.1101/2020.12.17.423316

**Authors:** Estefanía Calvo Alvarez, Serge Bonnefoy, Audrey Salles, Fiona E. Benson, Paul G. McKean, Philippe Bastin, Brice Rotureau

## Abstract

The single flagellum of African trypanosomes is essential in multiple aspects of the parasite development. The FLAgellar Member 8 protein (FLAM8), localised to the tip of the flagellum in cultured insect forms, was identified as a marker of the locking event that controls flagellum length. Here, we investigated whether FLAM8 could also reflect the flagellum maturation state in other stages. We observed that FLAM8 distribution extended along the entire flagellar cytoskeleton in mammalian infective forms. Then, a rapid FLAM8 concentration to the distal tip occurs during differentiation into early insect forms, illustrating for the first time the remodeling of an existing flagellum in trypanosomes. In the tsetse cardia, FLAM8 further localizes to the entire length of the new flagellum during an asymmetric division. Strikingly, in parasites dividing in the tsetse midgut and in the salivary glands, the amount and distribution of FLAM8 in the new flagellum was seen to predict the daughter cell fate. We propose and discuss how FLAM8 could be considered as a meta-marker of the flagellum stage and maturation state in trypanosomes.

**Summary statement:** The trypanosome protein FLAM8 displays a dynamic and stage-specific distribution during the entire parasite cycle, representing a novel marker of the flagellum stage and maturation state.

## Introduction

*Trypanosoma brucei* is an extracellular parasite responsible for African trypanosomiases, also known as sleeping sickness in humans and nagana in cattle. African trypanosomes are blood and tissue-dwelling protists transmitted to their mammalian hosts by the bite of the blood-feeding tsetse fly *(Glossina* genus) in sub-Saharan Africa. They are subjected to a complex developmental cycle highly organized in space and time, and characterized by the existence of multiple stages alternating between the two hosts (reviewed in (Rotureau and Van Den Abbeele, 2013)).

The trypanosome flagellum is an essential organelle anchored along the surface of the cell body and present in all stages of its development (Rotureau et al., 2011). It is essential for parasite viability (Broadhead et al., 2006), cell division and morphogenesis (Kohl et al., 2003), attachment to the tsetse salivary glands (Tetley and Vickerman, 1985) and motility (Rotureau et al., 2014b; Shimogawa et al., 2018), but it also possibly contributes to sensory functions and interactions with the microenvironment (reviews in (Roditi et al., 2016; Rotureau et al., 2009)). The trypanosome flagellum is composed of a canonical axoneme containing nine doublet microtubules and a central pair of singlet microtubules, associated with a paraflagellar rod (PFR) and surrounded by a specialized membrane (Langousis and Hill, 2014). This single flagellum exits the cytoplasm from the flagellar pocket, a specialized membrane invagination (Field and Carrington, 2009). It remains attached along most of its length to the cell body by the flagellum attachment zone with the exception of its distal end that extends to a short distance beyond the cell body (Taylor and Godfrey, 1969).

Unlike the microtubule-based core and the proximal basal body, our understanding of the molecular composition, specific structures and functions of the distal tip region is still limited, due mainly to its small dimensions and its continuity with other flagellar structural components. During flagellum construction, incorporation of axonemal precursors and PFR subunits only takes place at the distal tip of the growing flagellum (Bastin et al., 1999; Sherwin et al., 1987), which exhibits a complex ultrastructure that differs between insect and mammalian forms (Hoog et al., 2014; Hughes et al., 2013). In addition, the IntraFlagellar Transport (IFT) machinery mediates delivery of nascent flagellum subunits towards the ciliary tip for assembly into the growing flagellum and back to its base (Buisson et al., 2013). At the tip, the canonical axoneme organization collapses as microtubules are attached together in a resistive structure that prevents microtubule sliding but permits bend formation, which results in protractive tip-to-base flagellar beating (Woolley et al., 2006). Several studies have investigated the presence of discrete proteins that localise to the distal tip in trypanosomes, suggesting the existence of a specific subdomain (Chan and Ersfeld, 2010; Liu et al., 2010; Saada et al., 2014; Varga et al., 2017; Woodward et al., 1995).

Our proteomic analysis of preserved flagella purified from the insect midgut procyclic forms of the parasite (PCF) identified a group of flagellar membrane and matrix proteins with unique patterns and dynamics (Subota et al., 2014). Amongst them, one protein termed FLAgellar Member 8 (FLAM8) is present only at the distal tip of the flagellum of PCF grown in culture. This large protein (3,075 amino acids) is progressively added to the new flagellum during its assembly (Subota et al., 2014) and requires IFT to be maintained at the distal tip (Fort et al., 2016). In PCF trypanosomes, FLAM8 concentrates at the tip of axonemal microtubules after detergent extraction, (Subota et al., 2014) demonstrating its strong association to the flagellum cytoskeleton and/or an association to another specific structural complexes linked to the flagellum tip. Prior to PCF cell division, FLAM8 distribution in the distal part of the new organelle reaches about one third of that in the old flagellum (Subota et al., 2014). After cytokinesis, the amount of FLAM8 further increases until the flagellum reaches its final length. Recently, a new model termed “grow and lock” described how the new flagellum elongates until a locking event fixes the final length in a timely defined manner (Bertiaux et al., 2018). This study identified FLAM8 as a marker of the locking event that controls flagellum length and defines a flagellum that has reached its maturity status.

The grow-and-lock model results from observations in PCF cells cycling in stable culture conditions and that produce the same type of progeny. However, within each host, trypanosomes have to face different microenvironments, which requires major morphological and metabolic adaptations, driven by the activation of specific gene expression programs, that are critical for life-cycle progression (MacGregor et al., 2012; Ooi and Bastin, 2013; Smith et al., 2017). These drastic changes are also true for flagella that evolve in length, position and shape (Ooi and Bastin, 2013; Rotureau et al., 2011) as well as in molecular composition (Oberholzer et al., 2011; Rotureau et al., 2012; Subota et al., 2014). We reasoned that FLAM8 location and functions could also evolve during the parasite cycle. Especially, the role of FLAM8 as a marker of maturity at the tip of the flagellum could differ among specific stages, and FLAM8 could participate in flagellum length control in different ways through the trypanosome development.

Here, we detail the differential FLAM8 localization profiles over both the cell and parasite cycles *in vivo* in mammalian and insect hosts. FLAM8 amount and distribution along the trypanosome flagellum are dynamic and stage-specific. The first FLAM8 redistribution during the mammalian to insect form differentiation in the tsetse midgut illustrates the remodeling of an existing flagellum, while the second redistribution takes place during the construction of the new short flagellum in long asymmetrically dividing epimastigotes of the cardia. FLAM8 appears as a marker of the flagellum maturation state that could predict the fate of procyclic trypomastigotes dividing in the midgut and attached epimastigotes dividing in the salivary glands.

## Results

### FLAM8 is localized along the entire flagellum in BSF

Two trypanosome stages can be found in the bloodstream of a mammalian host: the dividing slender (SL) bloodstream form (BSF) and the non-proliferative stumpy (ST) form pre-adapted for transmission to the insect vector. To study the subcellular localization of FLAM8 in BSF trypanosomes, an anti-GFP antibody was used for immunostaining in combination with the mAb25 axonemal marker in parasites expressing FLAM8 coupled to EYFP at the N-terminal end. In contrast to PCF cells, FLAM8 was not only present at the distal tip of the flagellum, but it was also distributed along the entire flagellum length, with an enrichment in the proximal half of the flagellum in BSF in culture (Fig. 1A). At the BSF flagellum base, FLAM8 was detected after the emergence of the axoneme from the transition zone but before the beginning of the paraflagellar rod, possibly at the level of the flagellar pocket collar, evidenced by the use of the flagellar transition zone component (FTZC) (Bringaud et al., 2000) and PFR markers in BSF parasites expressing EYFP::FLAM8 (Fig. 1B).

**Figure 1.**
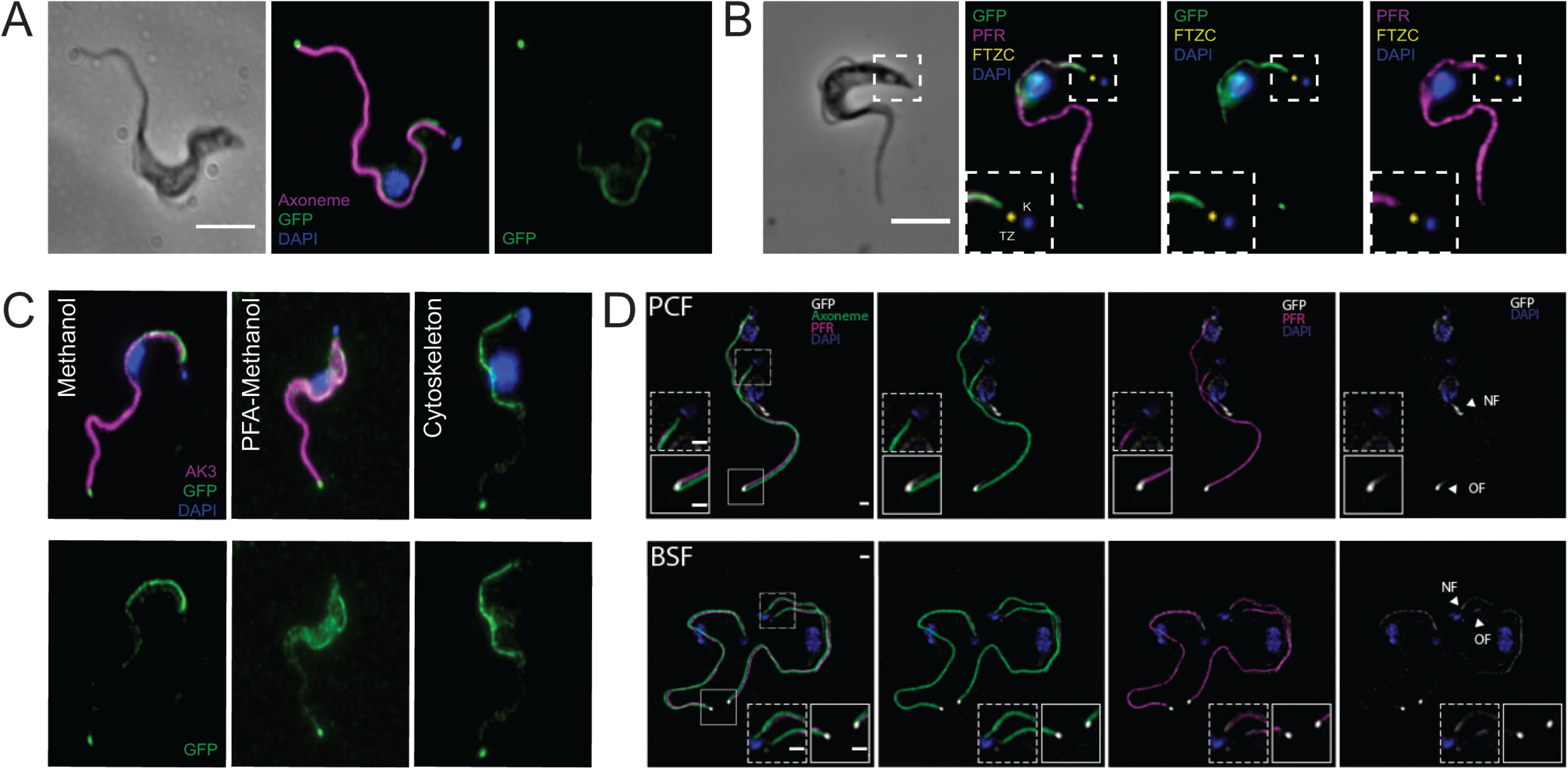
FLAM8 is differentially distributed along the entire flagellum in mammalian forms. **A)** Immunofluorescence on methanol-fixed AnTat EYFP::FLAM8 BSF in culture, using the anti-GFP (green), the anti-axoneme marker mAb25 (magenta), DAPI staining of DNA content (blue). Scale bar is 5 μm. **B)** IFA on methanol-fixed AnTat EYFP::FLAM8 BSF showing that at the base of the flagellum FLAM8 (anti-GFP signal in green) do not co-localize with the transition zone component FTZC (in yellow) and initiates before the starting of the PFR (in magenta). Scale bar is 5 μm. **C)** FLAM8 is not localized to the flagellum membrane. IFA on methanol-fixed AnTat EYFP::FLAM8 BSF (left panels), paraformaldehyde-fixed (middle panels) and cytoskeletons (right panels) using the anti-GFP antibody (green) and an antiserum against the arginine kinase of *T. cruzi* (magenta), DAPI staining of DNA content (blue). Arginine kinase (AK) signal, which is localized to the flagellum membrane, was lost after detergent extraction (right panels). Scale bar is 5 μm. **D)** Structured Illumination Microscopy images (SIM) of FLAM8 (anti-GFP antibody in white) with flagellar components, paraflagellar rod (PFR, magenta) and axoneme (green) showed in pleomorphic EYFP::FLAM8 PCF and BSF parasites (upper and lower panels, respectively). DAPI staining of DNA content appears in blue. Scale bars represent 1 μm. Images: white arrowheads show the old (OF) and the new flagellum (NF).

*Trypanosoma brucei* arginine kinase 3 (AK3) is localized to the entire flagellum membrane in all stages of parasite development in the tsetse fly as well as in BSF parasites (Ooi et al., 2015), representing a robust flagellum membrane marker. To assess whether FLAM8 could be associated with the BSF flagellum membrane, an antiserum raised against the arginine kinase of *Trypanosoma cruzi* (anti-*Tc*AK) (Pereira et al., 2002; Subota et al., 2014), was used for immunofluorescence assay in combination with the anti-GFP antibody in parasites expressing EYFP::FLAM8 either on methanol-fixed or paraformaldehyde-fixed whole cells, or detergent-extracted cytoskeletons. Distinct fluorescent signals were observed for FLAM8 and AK3 in the flagellum of parasites fixed in methanol and PFA-fixed (Fig. 1C, left and middle panels). This absence of overlap suggests that FLAM8 is not associated with the flagellum membrane. This was confirmed by the presence of FLAM8 at the proximal part of the flagellum and the distal tip after detergent extraction, whereas membrane removal led to a complete loss of the anti-AK3 signal (Fig. 1C, right panels). In order to define more closely the localization of FLAM8 to the flagellum cytoskeleton, Structured Illumination Microscopy (SIM) was employed with anti-GFP for FLAM8 staining, and markers for the axoneme and the paraflagellar rod (Kohl et al., 1999) in both EYFP::FLAM8 PCF and SL BSF (Fig. 1D). In PCF, the triple staining revealed that FLAM8 is exclusively located at the distal end of the flagellum in growing and mature flagella (Fig. 1D, upper panels). In contrast, bloodstream parasites exhibit a specific FLAM8 proximal localization that starts at the base of the flagellum and extends towards the half of the flagellum length in both old and new flagella (Fig. 1D, lower panels). At this point, the signal diminishes until reaching the distal tip of the cilium, where a bright FLAM8 accumulation is evidenced (Fig. 1D, lower panels). In both PCF and BSF, and in both mature and growing flagella, FLAM8 is distinct from, but resides in the vicinity of the axoneme and the PFR (Fig. 1D, upper and lower panels). This was observed in multiple cells and after calibration of the SIM settings with fluorescent microspheres to prevent any pixel shift. At the distal end, FLAM8 signal appears more extended and diffuse in PCF compared to the rounded and more defined shape observed in BSF. In addition, in the proximal region of the BSF flagellum only, SIM images confirm that FLAM8 signal initiates after the axoneme but before the PFR.

### FLAM8 is a marker of flagellum maturation in BSF

African trypanosomes possess a nucleus (N) and a single mitochondrion whose genetic material is condensed in a structure named kinetoplast (K) and that is physically linked to the basal body of the flagellum (Robinson and Gull, 1991). The progression through the cell cycle can be monitored by staining DNA content as parasites with one kinetoplast and one nucleus (1K1N) are in the G1/S phase, those with two kinetoplasts and one nucleus (2K1N) are in G2/M, and cells with two kinetoplasts and two nuclei (2K2N) are about to undergo cytokinesis (Sherwin and Gull, 1989; Woodward and Gull, 1990). FLAM8 was described as a molecular marker for flagellum maturation in cultured PCF as a maximum amount of FLAM8 is found in flagella that do not elongate anymore after having reached their mature state (Bertiaux et al., 2018). This led us to explore whether FLAM8 would have a similar behaviour in BSF trypanosomes *in vitro* and *in vivo* (Fig. 2 and S2). In order to perform quantitative analyses in flagella, both at the distal tip and at the proximal region, we used BSF trypanosomes expressing EYFP::FLAM8. The portion of the proximal region containing FLAM8 was measured in mature and growing flagella following a double immunofluorescence assay with an anti-GFP and the mAb25 axonemal marker for measurements. In 1K1N parasites, FLAM8 was observed in the first 13.5 μm ± 2.0 μm of the proximal region, representing 52% of the flagellum length, as well as at the distal tip of the mature flagellum (Fig. 2A, left panels and 2B). In 1K1N parasites containing a short new flagellum and in 2K1N trypanosomes undergoing mitosis, the presence of FLAM8 was restricted to the first 5.9 μm ± 3.2 μm and 11.6 μm ± 3.1 μm of the proximal region of the new flagellum, respectively (Fig. 2A, middle panels), which corresponded to 71% and 75% of the length of the growing flagellum, respectively (Fig. 2B). The FLAM8 distribution further extended up to 15.1 μm ± 3.8 μm in length in the new flagellum of 2K2N cells (Fig. 2A, right panels and 2B). At this point, the new flagellum has reached about 80% of its final mature length (not shown), implying that FLAM8 should still accumulate and/or re-locate along the full flagellum length after cell division until the growing flagellum has reached a definitive length and before the cell re-enters a new cell cycle. There was a linear correlation between the length of FLAM8 distribution in the proximal region and the total length of the axoneme at all steps of the cell cycle, except in the new flagella of 2K2N cells, where a less defined pattern was observed (Fig. 2C).

**Figure 2.**
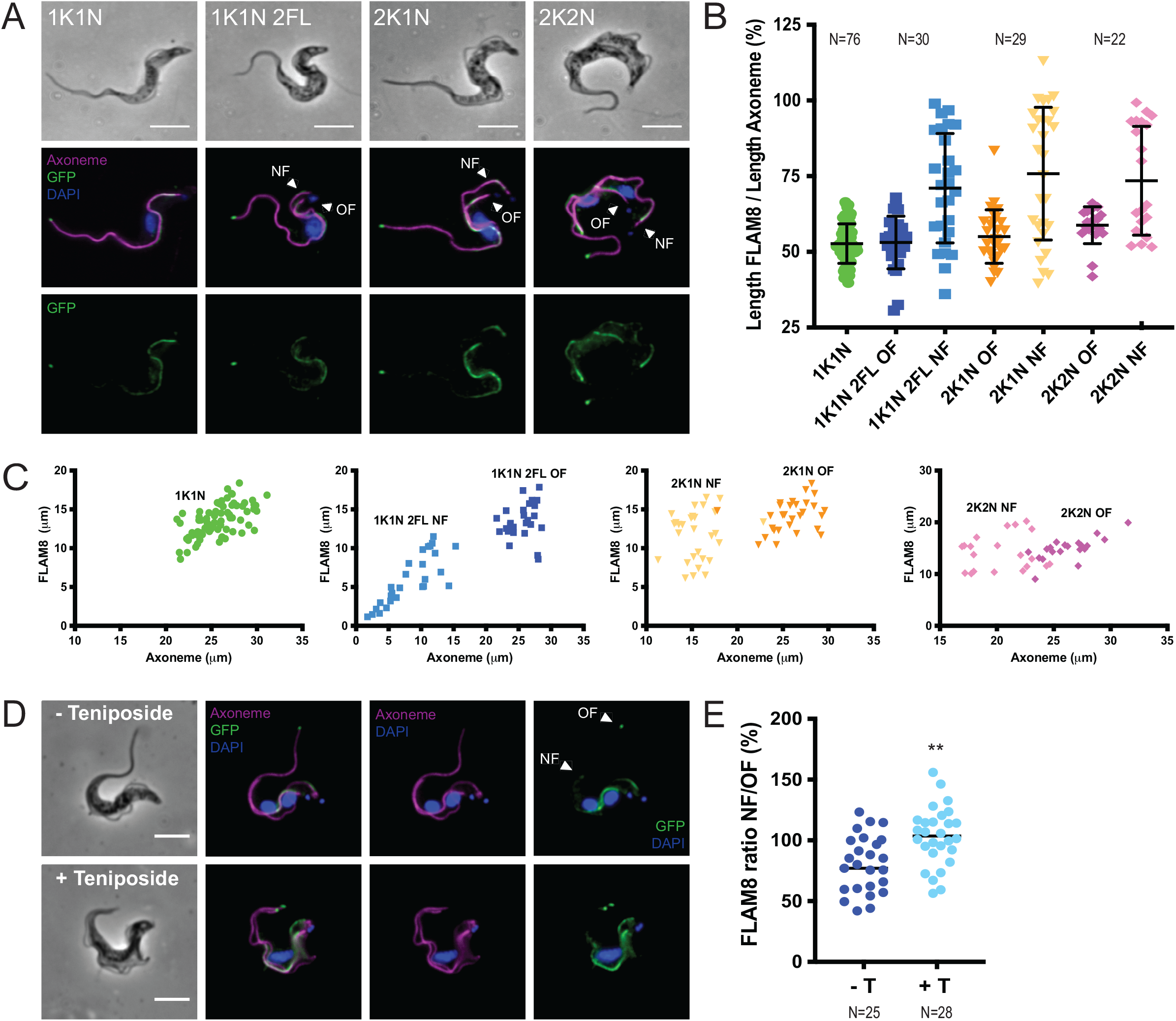
FLAM8 is also a marker of the flagellum maturation in mammalian infective forms. **A)** Representative immunofluorescence images on AnTat EYFP::FLAM8 BSF in culture showing distinct cell-cycle stages: 1K1N, 1K1N with a new short flagellum (1K1N 2FL), 2K1N and 2K2N parasites. The axoneme (mAb25 antibody in magenta) shows the full length of the flagellum; FLAM8 was labelled using the anti-GFP antibody (green), and DAPI staining of DNA content appears in blue. **B)** Dot plot showing the quantification of FLAM8 intensity present at the proximal region in 1K1N, 1K1N 2FL, 2K1N and 2K2N BSF trypanosomes, compared with the total length of the flagellum measured by the use of the axoneme marker mAb25. The number of cells used for the quantification (N) is indicated. **C)** Linear correlation between the FLAM8 signal and the length of the axoneme in 1K1N, 1K1N 2FL, 2K1N and 2K2N trypanosomes. The number of cells used for the quantification (N) is indicated above the graph in (B). **D)** IFA on methanol-fixed AnTat EYFP::FLAM8 parasites left untreated (top panels) or treated for 6 h with teniposide (bottom panels) stained with anti-GFP (green), mAb25 antibody (magenta) and DAPI labelling DNA (blue). **E)** Ratio between the FLAM8 fluorescent signal at the tip in the new and old flagellum in cells treated or non treated with teniposide for 6 h. Statistically significant differences are indicated with two stars (p < 0.005), and the number of cells used for quantification is indicated below the graph. Images: white arrowheads show the old (OF) and the new flagellum (NF). Scale bar is 5 μm.

To investigate whether FLAM8 distribution is dynamic and cell-cycle dependent in BSF as it is the case in PCF, cell division was inhibited by treating pleomorphic EYFP::FLAM8 BSF cells with 10 μM teniposide, a topoisomerase II inhibitor that interferes with mitochondrial DNA segregation but neither with basal body duplication nor with flagellum elongation (Robinson and Gull, 1991). In untreated cells, FLAM8 was abundant at the tip and in the proximal region of the old flagellum and present in lower amounts in the new flagellum (Fig. 2D, upper panels). By contrast, in cells treated with teniposide for 6 h (just under the cell doubling time), the new flagellum had elongated further and the FLAM8 signal at the distal region was brighter and looked similar to that at the tip of the mature flagellum (Fig. 2D, bottom panels). Unfortunately, the length of both old and new flagella could not be quantified due to a frequent overlapping between them in most cells after teniposide treatment. Next, regions of interest were defined around the tip of the new and old flagella, and the total amount of fluorescence was quantified. In untreated cells at an advanced stage of the cell cycle (2K2N), the ratio of the total FLAM8 fluorescence intensity between the new and the old flagella was 80% (Fig. 2E). However, in cells treated with teniposide, this ratio significantly increased up to 100% (Fig. 2E), demonstrating that a maximum amount of FLAM8 marker is found in flagella that have reached their final length in BSF trypanosomes as well. This confirms that FLAM8 can be considered as a flagellum maturation marker both in PCF and BSF trypanosomes.

### FLAM8 redistribution occurs in the existing flagellum during BSF to PCF differentiation

Several hours after the ingestion of a bloodmeal, ST differentiate into PCF parasites in the posterior midgut of the tsetse fly where they start to proliferate again while the digestion occurs. Since FLAM8 presents distinct distributions in BSF and PCF, we investigated how this transition could occur (Fig. 3). FLAM8 could be redistributed in non-dividing ST, or alternatively during either the SL to ST or the ST to PCF differentiation. In order to get insights into this remodeling, blood was sampled from the first peak of parasitemia of mice infected with trypanosomes and the redistribution of FLAM8 was investigated. The presence of stumpy cells was assessed by using the anti-PAD1 antibody marker that targets a unique surface transporter (Dean et al., 2009). The concomitant use of the anti-FLAM8 antibody showed that FLAM8 distributed along the full-length flagellum while exhibiting a bright signal at the distal tip in freshly isolated ST cells, as in SL trypanosomes (Fig. 3A, left panels). Then, the differentiation of these ST parasites into PCF was induced *in vitro* by a temperature shift from 37°C to 27°C coupled to the addition of cis-aconitate to the medium (Engstler and Boshart, 2004; Szoor et al., 2010). The emergence of PCF cells was assessed by using an anti-EP-procyclin marker antibody labelling only the procyclin surface coat (Acosta-Serrano et al., 2001). The expression of procyclin at the parasite surface was observed as soon as 2 h after differentiation (Fig. 3A, 2h-middle panels). This was concomitant with the FLAM8 enrichment to the distal tip, especially in cells with an intermediate morphology between BSF and PCF (Fig. 3A: 2h, 4h and 12h-middle panels). This remodeling culminated with the disappearance of the FLAM8 proximal signal and its concentration solely at the flagellum tip 24 h post-differentiation (Fig. 3A, right 24h-panels). The quantification of the FLAM8 signal at the distal tip in SL and ST BSF as compared to early PCF demonstrated the massive accumulation of the protein at the flagellum tip in PCF parasites (Fig. 3B). This accumulation of FLAM8 at the distal tip was also confirmed by repeating the same experiment with *in vitro* induced stumpy form parasites (Fig. S3, A-B). These spectacular modifications of FLAM8 location during ST BSF to PCF differentiation illustrates the remodeling of an existing flagellum in a non-dividing cell.

**Figure 3.**
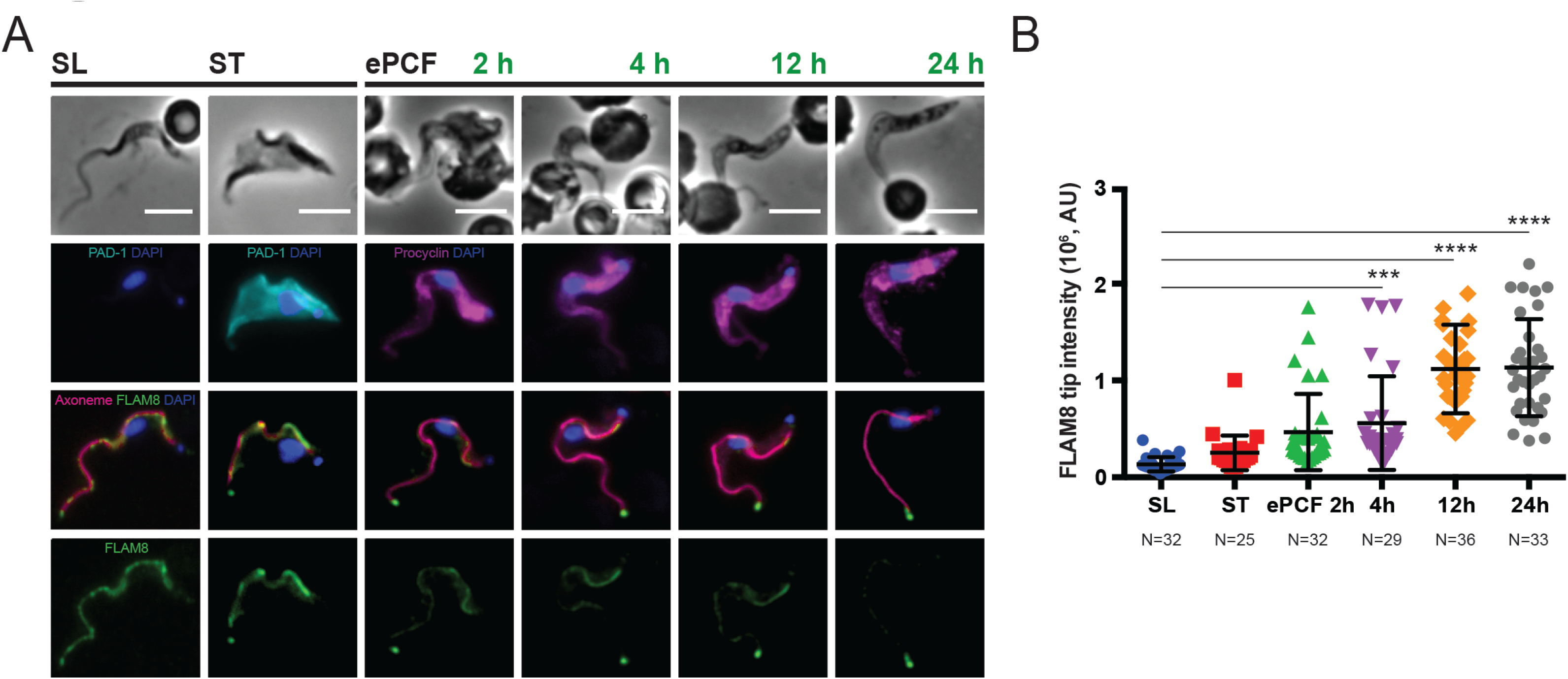
FLAM8 concentration to the tip reflects the remodeling of an existing flagellum during BSF to PCF differentiation. **A)** Representative images of AnTat 1.1E SL and ST parasites isolated from one infected BALB/c mouse and undergoing *ex vivo* differentiation from non-dividing stumpy forms (ST, left panels) into early procyclic trypanosomes at 27°C (ePCF, panels named 2h, 4h, 12h and 24h). Immunofluorescence images of methanol-fixed parasites stained with anti-FLAM8 antibody (green), anti-axonemal mAb25 (pink), and antibodies against specific surface markers: anti-PAD1 to stain ST forms (in cyan), and anti-procyclin labelling early PCF parasites (in magenta). DAPI staining appears in blue. Scale bar is 5 μm. **B)** Quantification of the FLAM8 intensity at the tip of the flagellum during *ex vivo* differentiation into insect procyclic forms. Statistically significant differences are indicated with three (p < 0.001) and four stars (p < 0.0001). AU: arbitrary units. The number of parasites used for quantification (N) is indicated below the graph.

### FLAM8 is a marker of cell fate after PCF division

During the first three days after their initial differentiation, PCF proliferate and eventually establish an infection in the posterior midgut (Gibson and Bailey, 2003). Later, only a subpopulation of these proliferative procyclic parasites progressively elongate and migrate to the anterior part of the midgut as long trypomastigotes, while the others stay in the posterior midgut and maintain the population (Van Den Abbeele et al., 1999). Assuming that PCF division results in the production of two daughter cells that are not equivalent (Abeywickrema et al., 2019), one could propose at least two hypotheses. First, only the daughter cell inheriting the new flagellum would be able to progress in the cyclical development (Ooi and Bastin, 2013). Second, alternatively or in addition, there could be two distinct types of PCF divisions: one division type leading to two daughter cells intended to further proliferate, and another division type producing one sibling adapted for progressing in the parasite cycle and a second sibling committed for another round of proliferation. In PCF parasites in culture that could virtually proliferate indefinitely, FLAM8 is progressively added to the new flagellum during its construction in G2/M phase and after cell division in G1 phase (Bertiaux et al., 2018; Subota et al., 2014). In order to confirm this *in vivo* and to explore the two hypotheses, we scrutinized the localization of FLAM8 in dividing PCF isolated from both the posterior and anterior regions of the tsetse midgut (Fig. 4) and treated them for IFA with the anti-FLAM8 antibody and the axonemal marker mAb25 (Dacheux et al., 2012). The FLAM8 signal intensity at the distal tip of old flagella (OF) in 1K1N, 2K1N or 2K2N cells was quantified and compared to that of new flagella (NF) in dividing 2K1N and 2K2N parasites, and the flagellum length was also measured (Fig. 4). In recently differentiated PCF isolated from the posterior midgut (3 days after the flies received an infective blood meal), a progressive accumulation of FLAM8 was observed in the growing flagellum of 2K1 N and 2K2N parasites (Fig. 4A). The total amount of FLAM8 at the tip of the new flagellum was significantly lower than that in the mature flagellum in 2K1 N cells (Fig. 4B). A similar yet non-significant trend was also evidenced in 2K2N cells (Fig. 4B-C). This demonstrates that in procyclic trypanosomes from the posterior midgut, FLAM8 addition to the distal tip occurs during the assembly of the new flagellum and that the protein reaches its maximum amount in the new flagellum after cell division, as observed in PCF in culture (Bertiaux et al., 2018; Subota et al., 2014). Nevertheless, a different situation was observed in late PCF extracted from the anterior midgut (Fig. 4D-F). Eleven days after ingestion of the blood meal, trypanosomes had migrated and colonized the anterior region, where they were found in high numbers. A region of interest was defined around the tip of all flagella in 1K1N, 2K1N and 2K2N cells from the AMG, and the total amount of fluorescence was quantified (Fig. 4E). Surprisingly, the amount of FLAM8 at the tip of the new flagellum was significantly higher than in the old flagellum in 2K2N trypanosomes (Fig. 4E). Actually, the amount of FLAM8 in the growing flagellum of a 2K2N PCF corresponded to that in the mature flagellum of a 1K1N PCF from the anterior midgut (Fig. 4E). These findings contrast with the opposite profile seen in PCF dividing in the posterior midgut (Fig. 4A) and with previous observations in cultured trypanosomes (Bertiaux et al., 2018; Subota et al., 2014). In parallel, the amount of FLAM8 in the old flagellum was significantly decreasing during cell division (Fig. 4E, compare red squares with green triangles). Furthermore, significant differences were observed when the ratios of FLAM8 intensities in the NF vs. OF of dividing 2K1N and 2K2N were compared in parasites coming from the posterior and the anterior midgut regions (Fig. 4C and 4F, respectively).

**Figure 4.**
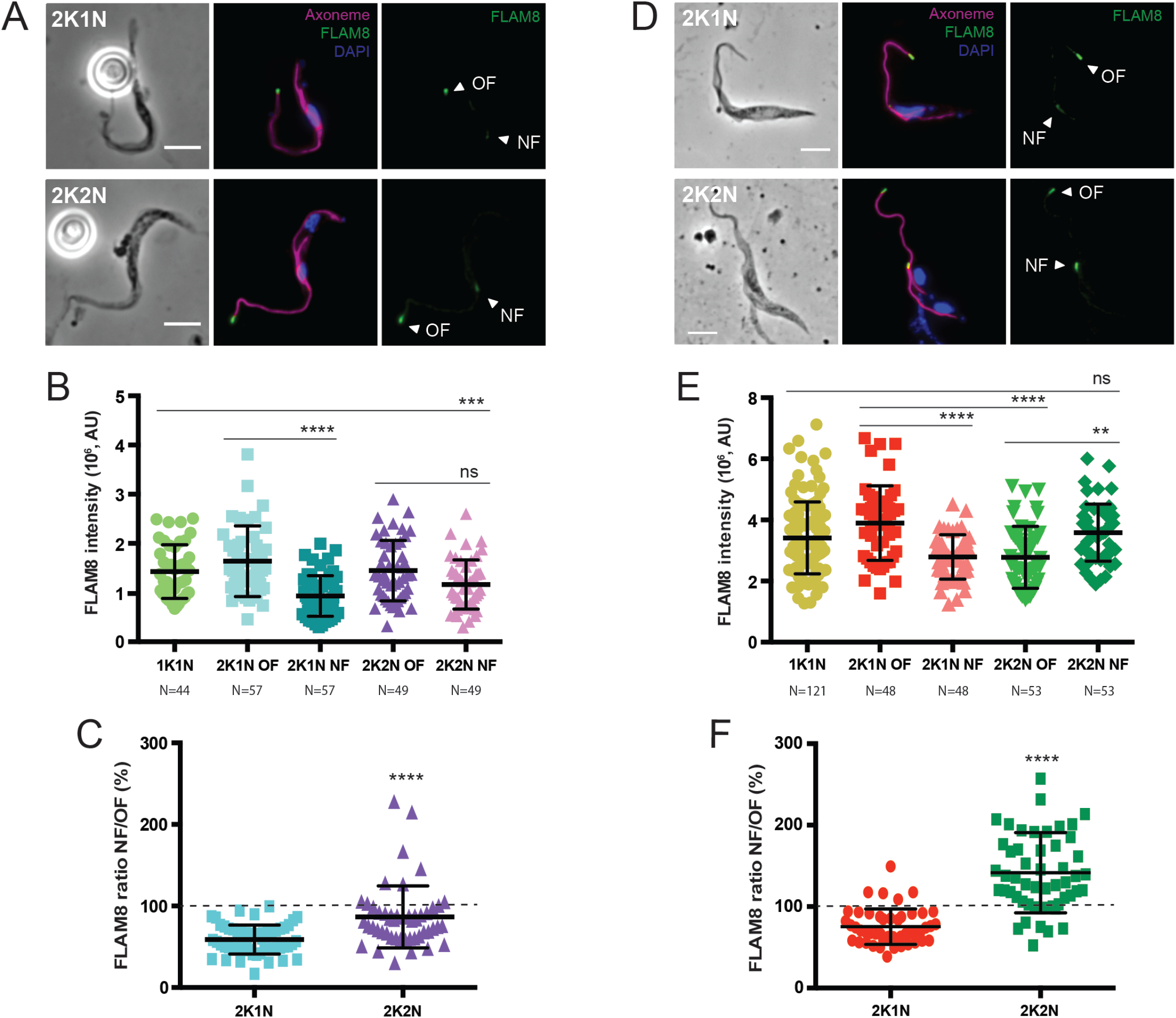
FLAM8 accumulates differently in dividing procyclics from posterior vs. anterior midgut. *Glossina m. morsitans* tsetse flies were fed with a blood meal containing pleomorphic AnTat 1.1E stumpy BSF parasites. The flies were dissected at distinct time points in order to isolate different MG regions and look for PCF trypomastigote stages. **A-C)** Infected tsetse flies were dissected at day 3 for posterior midgut (PMG) region isolation. **A)** Representative immunofluorescence images of methanol-fixed 2K1N (upper panels) and 2K2N (bottom panels) dividing parasites stained with anti-FLAM8 antibody (green), anti-axoneme marker mAb25 (magenta) and DAPI staining (blue). Scale bar is 5 μm. **B)** Quantification of FLAM8 intensity at the flagellum tip of 1K1N parasites and both old (OF) and new flagella (NF) of dividing 2K1N and 2K2N trypanosomes. Statistically significant differences are indicated with three (p < 0.001) and four stars (p < 0.0001); ns: not significant. **C)** Ratio between FLAM8 intensity in the new flagellum (NF) and the old flagellum (OF) in 2K1N and 2K2N trypanosomes. Statistically significant differences are indicated with four stars (p < 0.0001). Black dotted line indicates signal at 100% ratio. **D-F)** Tsetse flies receiving an infective blood meal were dissected after 11 days for anterior midgut (AMG) isolation. **D)** Representative immunofluorescence images of methanol-fixed 2K1N (upper panels) and 2K2N (bottom panels) procyclic cells showing anti-FLAM8 (green), mAb25 signals (magenta) and DAPI staining (blue). Scale bar is 5 μm. **E)** Dot plot showing FLAM8 signal quantification at the flagellum tip of 1K1N parasites and 2K1N and 2K2N procyclic cells at both mature (OF) and growing flagella (NF). Statistically significant differences are indicated with two (p < 0.01) and four stars (p < 0.0001). **F)** Ratio between FLAM8 intensity in the new flagellum (NF) and the old flagellum (OF) in 2K1N and 2K2N trypanosomes. Statistically significant differences are indicated with four stars (p < 0.0001). Black dotted line indicates signal at 100% ratio. In (B) and (E), the number of parasites used for quantification (N) is indicated below the graphs. Images: white arrowheads show the old (OF) and the new flagellum (NF).

Then, we reasoned that these differences in FLAM8 distribution patterns between dividing PCF in posterior *versus* anterior midgut could be related to the fate of their respective daughter cells. To confirm this possibility, FLAM8 localisation profiles were scrutinized in all trypomastigote stages from the midgut and cardia 14 days after ingestion (Fig. 5). FLAM8 signal at the distal tip appeared to increase with the parasite development, therefore, the total amount of fluorescence at the tip was subsequently quantified. Since the parasite migratory movement towards the cardia is accompanied by the progression through distinct developmental stages and concomitant with significant changes in flagellar length (Ooi and Bastin, 2013), this latter parameter was also measured. All trypomastigote parasites were characterized by i) the length of their flagellum assessed by the axonemal marker mAb25, and ii) their nuclear morphology ranging from a typical rounded shape in PCF from posterior midgut to an elongated conformation in long trypomastigotes (LT) from the anterior midgut and the cardia (Rotureau et al., 2011; Sharma et al., 2008). Interestingly, the amount of FLAM8 at the tip of long trypomastigotes in the cardia was significantly higher than that in PCF from the posterior midgut, and FLAM8 signal appeared more elongated compared to the condensed material found in PCF (Fig. 5A-B). Furthermore, the expression of FLAM8 correlated with flagellum length (Fig. 5C), increasing during trypomastigote parasite progression into the development from the posterior midgut to the cardia.

**Figure 5.**
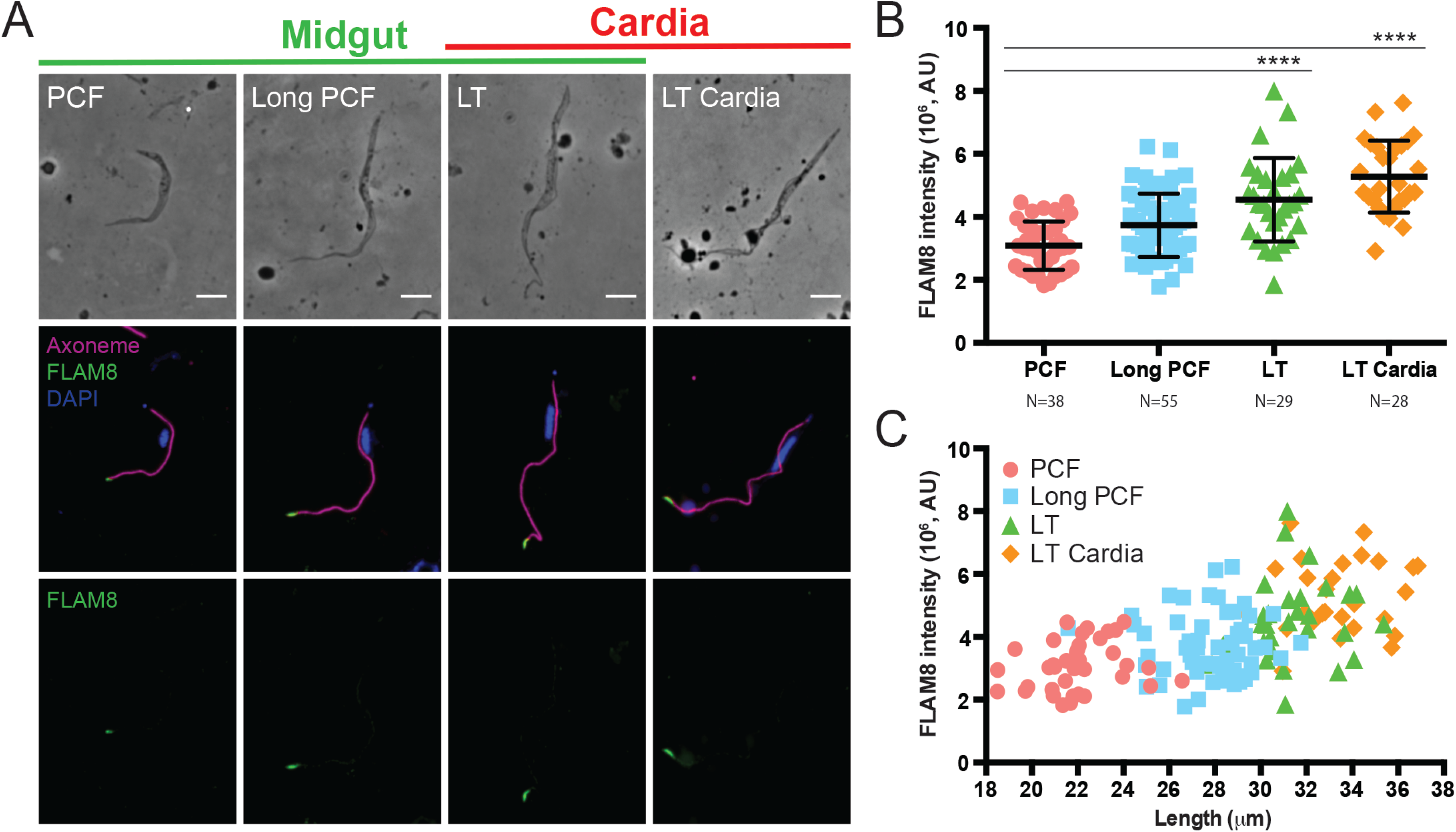
FLAM8 distribution at the flagellum tip extends with the elongation of the flagellum in long trypomastigote parasites in the midgut. Tsetse flies were fed with freshly-differentiated AnTat 1.1E stumpy BSF parasites and after 14 days the entire digestive tract including the cardia/proventriculus was isolated for immunofluorescence analyses and further quantification. **A)** Selected images of methanol-fixed procyclics (PCF), long PCF, long trypomastigotes (LT) and LT from the cardia stained with mAb25 axonemal marker (magenta), anti-FLAM8 antibody (green) and DAPI for DNA content (blue). Upper line: midgut stages (green) vs. trypomastigotes found in the cardia (red). **B)** Dot plot showing the FLAM8 intensity at the tip of PCF parasites, long PCF and LT recovered from the midgut and the cardia. Stastistically significant differences are indicated with four stars (p < 0.0001). The number of parasites used for quantification (N) is indicated below the graph. **C)** The mAb25 axonemal signal was used for flagellum length quantification and further correlation with the FLAM8 amount at the tip of specific parasite stages.

### FLAM8 distribution along the trypanosome flagellum is stage-specific

In the tsetse cardia, long dividing epimastigote trypanosomes maintained a similar FLAM8 localization (Fig. 6A). These cells divide asymmetrically to produce a long and a short epimastigote parasites (Van den Abbeele et al., 1999; Sharma et al., 2008). While long epimastigotes showed a FLAM8 localization concentrated at the tip, short epimastigote cells (the shortest morphological form identified with a flagellum length about 3-4 μm), exhibited a distinct distribution of FLAM8 along the entire flagellum length (Fig. 6A). Indeed, short epimastigote trypanosomes are assumed to be transported by the highly motile long epimastigotes into the lumen of the salivary gland, where they will attach to the epithelium and differentiate into attached epimastigote forms. The pattern observed in short epimastigotes was also observed in salivary gland parasites, i.e. in both attached epimastigote trypanosomes and infective metacyclic trypomastigote cells (Fig. 6A). A quantitative analysis of the fluorescence was performed in all stages by calculating the ratio between the integrated intensity of the FLAM8 signal in the flagellum and the flagellum length (Fig. 6B and C), and two intensity analyses were performed: one based on the FLAM8 signal found at the distal tip by drawing a ROI with a fixed size for all parasite stages (Fig. 6B), and one along the entire flagellum length by using the mAb25 signal labelling the axoneme (Fig. 6C). In both quantitative analyses, the shortest flagellum corresponding to the new flagellum of the dividing epimastigotes and the flagellum of the short epimastigotes presented the highest ratios, compatible with the redistribution / dilution of FLAM8 to the entire length of the flagellum as early as in short epimastigote parasites from the cardia.

**Figure 6.**
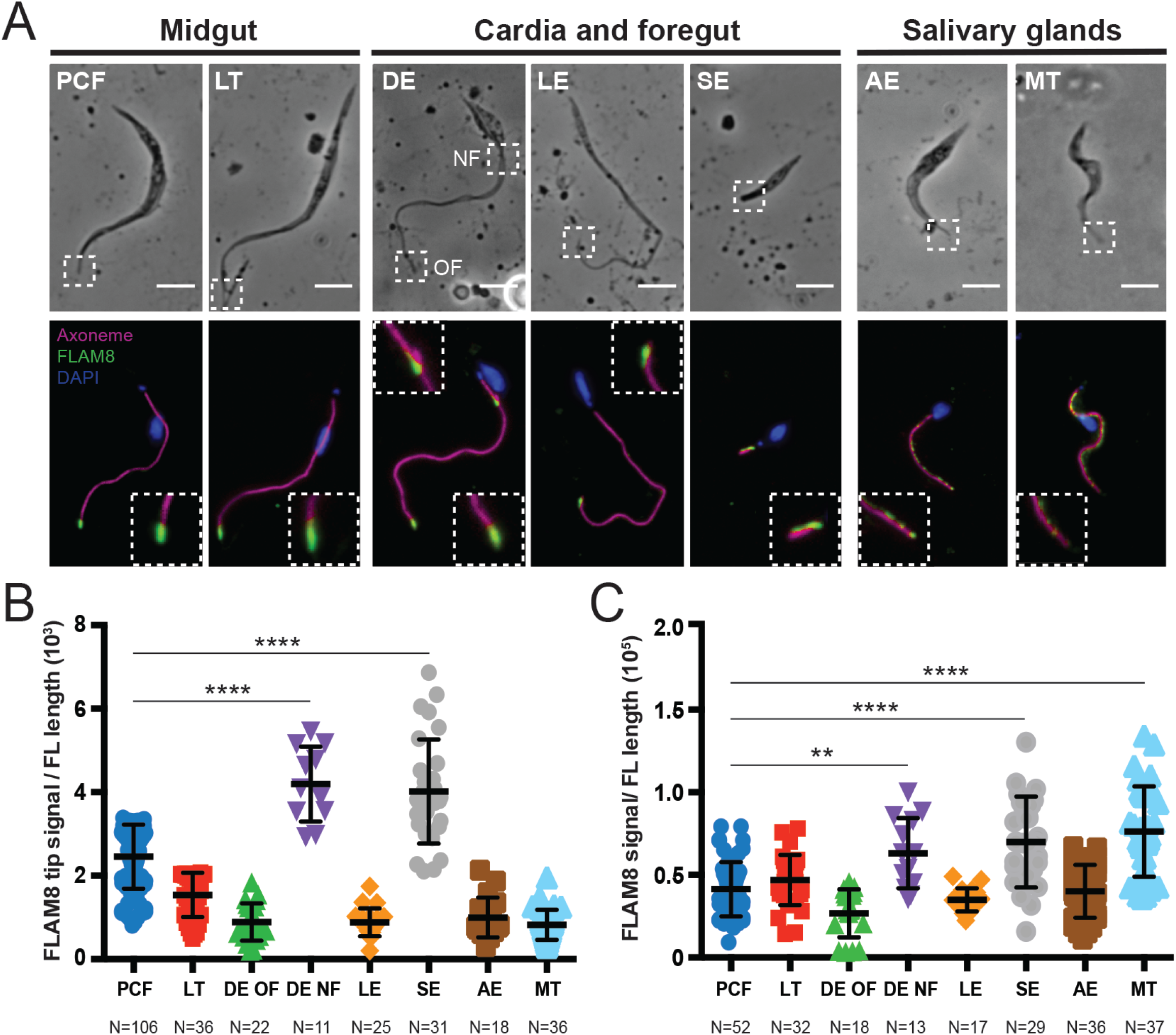
FLAM8 is redistributed along the trypanosome flagellum twice during the parasite cycle in the insect vector. **A)** AnTat 1.1E stumpy parasites were used to infect *Glossina m. morsitans* tsetse flies in order to collect fly tissues (midgut, cardia and salivary glands), and look for all developmental stages after 30 days of infection. Immunofluorescence images using anti-FLAM8 antibody (green), anti-axonemal marker mAb25 (magenta) and DAPI staining (blue). White dashed-line boxes show a two-fold magnification of the distal tip in all parasite stages. Images: PCF, procyclic; LT, long trypomastigote; DE, dividing epimastigote; LE, long epimastigote; SE, short epimastigote; AE, attached epimastigote; MT, metacyclic. Scale bar is 5 μm. **B, C)** The ratio of intensity of FLAM8 fluorescence at the distal tip (B) and along the entire flagellum (C) related to flagellum length was quantified in all trypanosome morphotypes found within the midgut, cardia and the salivary glands of infected tsetse flies. The number of cells used for the quantification (N) is indicated below the graphs. Stastistically significant differences are indicated with two (p < 0.01) and four stars (p < 0.0001).

Then, in the salivary glands, a continuous production of infective metacyclic parasites is ensured through an asymmetric division that results in two distinct daughter cells, one inheriting the old flagellum that displays the epimastigote configuration and the one inheriting the new flagellum which adopts the trypomastigote configuration (Rotureau et al., 2012). In order to investigate in more depth the dynamics of FLAM8 redistribution in salivary gland parasites undergoing metacyclogenesis, transition stages between epimastigote and trypomastigote, i.e. Epi-Trypo dividing cells, were analyzed (Fig. 7). Interestingly, during the asymmetric division in 2K1 N Epi-Trypo cells, a higher amount of FLAM8 was found along the growing new flagellum (Fig. 7A-B), a significant enrichment that was maintained in pre-metacyclic cells and even more pronounced in infective metacyclic trypomastigotes (Fig. 7A-B).

**Figure 7.**
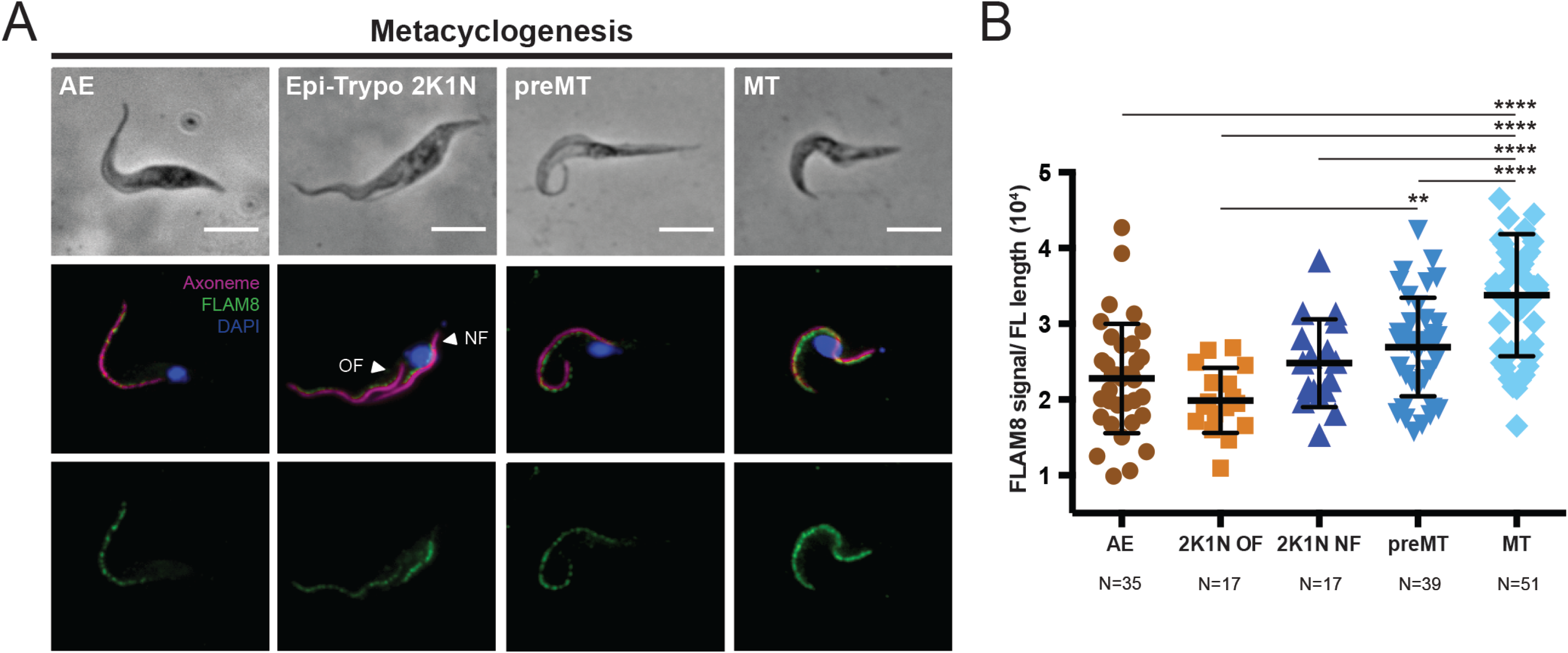
FLAM8 is differentially distributed during the asymmetric division in the salivary glands. **A)** Representative images of Epi-Trypo transition in the salivary glands. Methanol-fixed AnTat 1.1E parasites were stained with anti-FLAM8 antibody (green), anti-axonemal marker mAb25 (magenta) and DAPI staining (blue). Images: AE, attached epimastigote; 2K1N: Epi-Trypo parasite during the asymmetric division; preMT: pre-metacyclic; MT: metacyclic. White arrowheads within the Epi-Trypo panel indicate old (OF) and new (NF) flagellum. Scale bar is 5 μm. **B)** Quantification of FLAM8 fluorescence intensity along the flagellum related to the length of the axoneme. The number of parasites considered for quantification (N) is indicated below the graph. Stastistically significant differences are indicated with two (p < 0.01) and four stars (p < 0.0001).

## Discussion

### 1. FLAM8 and the flagellum tip

FLAM8 was first detected at the distal tip of PCF trypanosomes in culture (Subota et al., 2014), an observation that has been confirmed in insect-derived PCF parasites in the present study. Only a small subset of flagellar proteins have also been exclusively localized to the distal end of the flagellum in PCF, suggesting the existence of a very specific microdomain (Chan and Ersfeld, 2010; Liu et al., 2010; Saada et al., 2014; Varga et al., 2017; Woodward et al., 1995). The repetition of PFAMB506 domains (Pfam ID PB000506), also found in 4 *T. brucei* flagellum attachment zone proteins as well as in a few other cytoskeletal associated proteins (Sunter et al., 2015), could be involved in the interactions of FLAM8 with cytoskeletal partners at the flagellum tip, albeit the function of this domain remains unknown. However, FLAM8 was not detected by immunoprecipitation in PCF flagellar tip extracts (Varga et al., 2017). As proposed in *Chlamydomonas* where HSP70 and EB1 have been localized to the distal tips of the flagella (Bloch and Johnson, 1995; Pedersen et al., 2006), FLAM8 was speculated to be involved in microtubule assembly, stability and / or dynamics at the distal tip of the axoneme. In addition, the redistribution of FLAM8 along the entire flagellum in BSF indicates that FLAM8 may not exclusively and permanently reside within such a distal tip location.

FLAM8 amount increases during flagellum construction and after cell division in PCF in culture (Subota et al., 2014), and it is maintained at the distal tip of the flagellum by IFT activity (Fort et al., 2016). Here, in both dividing BSF and PCF isolated from infected blood and tsetse midguts respectively, we confirmed that FLAM8 is progressively added to the tip of the new flagellum during its assembly. Other flagellar tip proteins display a similar pattern: the Axonemal Capping protein (ACS1), whose amount in the new flagellum is length-dependent in PCF (Varga et al., 2017), and two insect stage-specific adenylate cyclases, ACP1 and ACP4 (Saada et al., 2014). FLAM8 proteins newly synthesized in the cytoplasm need to transit through the transition zone that acts as a gate between the flagellum and the cell body compartments (Reiter et al., 2012). Although incorporation of FLAM8 in the new flagellum in dividing PCFs in the posterior midgut is likely to result from *de novo* synthesis of fresh FLAM8 material brought from the base to the tip by IFT, it could conceivably be different in the case of dividing PCFs in the AMG. Possibly in addition to *de novo* synthesis, the concomitant FLAM8 amount increase in the new flagellum and decrease in the old one during cell division in PCFs from anterior midgut could also result, at least partially, from a recycling of FLAM8 from the mature flagellum to the new flagellum. Some of the FLAM8 proteins present at the tip of the old flagellum could be transferred to the new flagellum by IFT successively via the IFT retrograde machinery in the old flagellum and the IFT anterograde machinery of the new flagellum, via the flagellar bases in the cytoplasm, although an exchange between the two flagella has not been proven so far. A variation in the number of FLAM8 binding sites at the tip of the old flagellum, with a free circulation of FLAM8 between the two flagella could also be envisaged.

### 2. FLAM8 as a flagellum maturation marker

Recently, the grow-and-lock model was proposed to explain how flagellum length could be regulated in procyclic trypanosomes in culture (Bertiaux et al., 2018). FLAM8 was identified as a marker of the locking event associated to the acquisition of the final flagellum length. Here, FLAM8 was also seen to progressively accumulate at the tip during the entire flagellum elongation process in SL BSF and the inhibition of cell division resulted in the production of BSF parasites bearing two long flagella with equal amount of FLAM8 at the distal tip, in contrast to dividing cells where FLAM8 signal in the new flagellum only reaches 80% of the amount present in the old one. These observations suggest that the locking event, whose exact molecular nature needs to be elucidated, is controlled by a general cellular process that is temporally regulated, and could possibly be involved in discriminating the progression through another cell cycle or a new round of differentiation.

In line with the proposal that PCF division results in the production of two daughter cells that are not equivalent (Abeywickrema et al., 2019), we reasoned that the discrepancy between the two FLAM8 distribution patterns observed between the two flagella of early dividing PCF from the posterior midgut (PMG) and late dividing PCF from the anterior midgut (AMG) could somehow reflect the distinct fate of their respective daughter cells bearing the new flagellum (Fig. 8). Early dividing PCF would keep on proliferating at a regular rate by producing daughter cells destined to further re-enter new cell division cycles in the PMG. In this context, FLAM8 is detected in lower amount in the new flagellum than in the old flagellum in 2K2N early PCF, resulting in the production of a daughter cell with a slighthly shorter flagellum that would further mature and elongate after cytokinesis, up to the same length as that of the old flagellum (Fig. 8), as proposed in the grow-and-lock model (Bertiaux et al., 2018). By contrast, part of the PCF population is assumed to elongate and to migrate to the tsetse cardia before differentiating into long asymetrically dividing epimastigotes (Lemos et al., 2020; Rotureau and Van Den Abbeele, 2013). Interestingly, we found that FLAM8 amount at the flagellum tip increases with the elongation of the flagellum during trypomastigote parasite progression from the PMG towards the anterior region and, finally, in the cardia. A second population of dividing PCF that would first migrate toward the AMG and start dividing in a different manner in order to produce the future long trypomastigotes that would further enter differentiation into the next stage was further characterised. In this case, late PCF found to divide in the AMG would do so in an asymmetrical manner, as assessed by the peculiar FLAM8 distribution in the two flagella: FLAM8 is detected in higher amounts in the new flagellum than in the old flagellum in 2K2N late PCF, resulting in the production of a daughter cell with a flagellum that would further mature and elongate after cytokinesis (Fig. 8). This situation could be compared to the differential distribution of calflagins in Epi-Trypo asymmetrically dividing cells in the salivary glands, resulting in the production of two distinct cells with different fates (Rotureau et al., 2012). Thus, we propose that the amount of FLAM8 at the tip of the new flagellum of 2K2N PCF may represent a potential marker of prediction for the future of the daughter cell that could become either a proliferative PCF in the posterior midgut or a long trypomastigote about to differentiate into epimastigote in the anterior midgut (Fig. 8).

**Figure 8.**
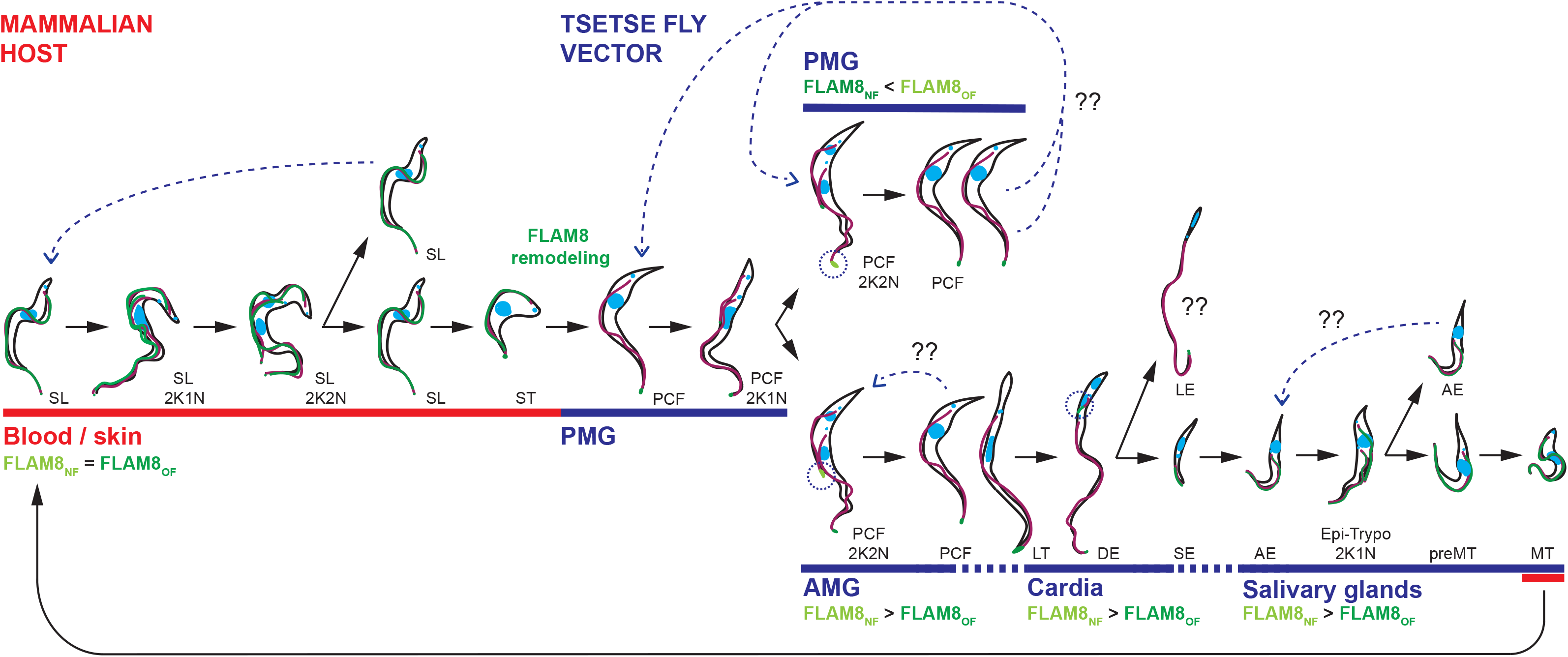
FLAM8 is a marker of the flagellum maturation state that predicts the fate of a trypanosome during the parasite cycle. In the mammalian host, proliferative slender (SL) parasites divide in order to colonize the intravascular and extravascular compartments. This symmetric division gives rise to one daughter cell that will undergo another round of division or that will stop proliferation and differentiate into arrested stumpy trypanosomes (ST). Both SL and ST trypomastigotes exhibit FLAM8 along the full flagellum length. During a blood meal on an infected host, a tsetse fly will ingest a pool of SL and ST cells, but only the latter will be capable of surviving in the new host environment. In the PMG, ST will rapidly transform into proliferative procyclic (PCF) trypomastigotes. This event will be characterized by the dramatic remodeling of FLAM8 distribution in the existing flagellum, since PCF cells display a FLAM8 concentrated only at the very distal tip of the flagellum. In addition, 2K2N PCF cells dividing in the posterior midgut (PMG) express a lower amount of FLAM8 in the growing flagella, associated with a post-division maturation of the NF in the daughter cell according to the grow-and-lock model. The daughter cell would then probably reenter another proliferation cycle in order to maintain the proliferative PCF population in the PMG. In contrast, dividing procyclic cells located in the anterior midgut (AMG) present a higher amount of FLAM8 in the NF before cytokinesis, suggesting that the daughter cell would proceed to the next developmental step and differentiate into nonproliferative long trypomastigotes (LT) by a drastic elongation of their flagella. These elongated trypomastigote parasites will subsequently enter the cardia in order to initiate a differentiation into epimastigote stages. During the asymmetric division performed by these long dividing epimastigotes (DE), two daughter cells are generated: a long epimastigote (LE), which is believed to die shortly after migration and/or cytokinesis and a short epimastigote (SE), that will further develop in the salivary glands. Unlike DE and LE, SE present FLAM8 along the entire length of the flagellum, evidencing a second redistribution of the protein *in vivo.* Once in the salivary glands, SE will differentiate into stages that will attach to the salivary epithelium, keeping FLAM8 along their entire flagellum. Interestingly, during the asymmetric division carried out by Epi-Trypo dividing AE, the daughter cell inheriting the NF exhibits a higher amount of FLAM8 as compared to the OF. This daugther cell will then differentiate into pre-metacyclic (preMT) parasites and, finally, into infective metacyclic trypomastigotes, with FLAM8 being present along the entire flagellum length. Legends: K, kinetoplast; N, nucleus; FLAM8_NF_, FLAM8 amount in the new flagellum; FLAM8_OF_, FLAM8 amount in the old flagellum.

### 3. Differential distribution of FLAM8 during the parasite cycle

The model presented in Figure 8 recapitulates all our observations and highlights that FLAM8 amount and distribution along the trypanosome flagellum are dynamic and stage-specific. A number of stage-specific proteins are well-characterized in African trypanosomes, especially surface antigens such as Variant Surface Glycoproteins, procyclin (Roditi et al., 1987) or *Brucei* Alanine-Rich Protein (Urwyler et al., 2007), but only a few stage-specific markers are known to be restricted to the flagellum. Members of the calflagin family are expressed during metacyclogenesis in attached epimastigote parasites asymmetrically dividing in the tsetse salivary glands. These membrane-associated proteins are progressively enriched exclusively in the new flagellum of the trypomastigote daughter cell that will further mature into an infective metacyclic form in the saliva (Rotureau et al., 2012). By contrast, here, by *ex vivo* differentiation of BSF into PCF parasites, we demonstrated that FLAM8 redistribution takes place in a few hours in 1K1N cells with a procyclic morphology. To our knowledge, this is the first evidence of a molecular remodeling occurring in the existing flagellum of a non-dividing trypanosome. This raises a number of questions regarding the mechanisms involved in this event and its impact on the flagellum length control. A quantitative phosphoproteomic comparison of BSF and PCF trypanosomes revealed differential protein phosphorylation levels among flagellar and RNA-binding proteins that are crucial for parasite differentiation (Urbaniak et al., 2013). This is the case of FLAM8, that presents differential phosphorylation between the two stages with 4 sites phosphorylated exclusively in PCF and a distinct one in BSF. This could be linked to its differential distribution in the two stages, hence possibly to distinct functions.

FLAM8 is then found concentrated at the tip of all trypomastigote stages in the midgut in amounts directly related to their flagellum length. However, it is detected along the entire length of the flagellum in short epimastigote cells (Fig. 8). This change of FLAM8 profile takes place during the construction of the new short flagellum in long asymmetrically dividing epimastigotes. At this step also, FLAM8 could be involved in flagellum length control, with a higher amount of FLAM8 being already produced in short epimastigotes to anticipate the upcoming rapid flagellum elongation necessary for a proper attachment to the salivary gland microvilli. Recent evidence has proved that trypanosome stages developing in the insect vector possess a total amount of IFT proteins that directly correlates with the length of the flagellum (Bertiaux et al., 2020). Interestingly, short epimastigotes were found as the sole parasite stage to exhibit a supplementary large amount of dynamic IFT material at the flagellar distal end (Bertiaux et al., 2020). In the salivary gland stages, FLAM8 is also a marker of the flagellum maturation state that appears to predict the fate of attached Epi-Trypo dividing cells, with an enrichment of FLAM8 in the flagellum in the trypomastigote configuration as compared to the epimastigote one (Fig. 8). This situation could also be compared to the differential distribution of calflagins in Epi-Trypo asymmetrically dividing cells (Rotureau et al., 2012).

In summary, we propose that FLAM8 could be considered as a meta-marker of the flagellum stage and maturation state in trypanosomes. FLAM8 distribution in trypanosomes is stage-specific and its remodeling seems to occur in two different situations: (i) in a new flagellum produced during cell divisions in PCF of the anterior midgut, in long epimastigotes of the cardia and in Epi-Trypo attached to the salivary glands, (2) or in the existing flagellum of non-dividing ST BSF differentiating into PCF in the midgut. Noteworthingly, this drastic remodeling of FLAM8 distribution in an existing flagellum is concomitant to major environmental changes for the parasites. Further research is needed to fully understand the roles of FLAM8 in the tsetse insect vector. In addition, understanding the potential protein interactions with other flagellum partners, especially at the distal tip, is another challenge that would help to unravel the essential roles of this fascinating and essential organelle.

## Materials and methods

### Strains, culture and *in vitro* differentiations

The AnTat 1.1E Paris pleomorphic clone of *Trypanosoma brucei brucei* was derived from a strain originally isolated from a bushbuck in Uganda in 1966 (Le Ray et al., 1977). Procyclic form trypanosomes (PCF) were grown at 27°C in SDM-79 medium (Brun and Schonenberger, 1979) supplemented with 10% (v/v) heat-inactivated foetal bovine serum (FBS), haemin (7.5 mg/ml) and 8 mM glycerol (SDMG). Bloodstream form trypanosomes (BSF) of the same strain were cultivated in HMI-11 medium supplemented with 10% (v/v) FBS (Hirumi and Hirumi, 1989) at 37°C in 5% CO_2_. For *in vitro* slender to stumpy BSF differentiation, we used 8-pCPT-2’-O-Me-5’-AMP, a nucleotide analog of 5’-AMP from BIOLOG Life Science Institute (Bremen, Germany). Briefly, 2×10^6^ pleomorphic AnTat 1.1E slender forms were incubated with 8-pCPT-2’-O-Me-5’-AMP (5 μM) for 48 h (Barquilla et al., 2012). Freshly differentiated stumpy form parasites were then centrifuged at 1,400 × g for 10 minutes in TDB (Trypanosome Dilution Buffer: 5 mM KCl, 80 mM NaCl, 1 mM MgSO_4_*7H_2_O, 20 mM Na_2_HPO_4_, 2 mM NaH_2_PO_4_, 20 mM glucose) and resuspended in SDMG medium with 10 mM glutathione for tsetse fly infection (MacLeod et al., 2007). For *in vitro* differentiation into PCF, freshly differentiated short stumpy form trypanosomes were transferred into SDM-79 medium supplemented with 10% (v/v) FBS, 6 mM cis-aconitate and 20 mM glycerol at 27°C (Czichos et al., 1986).

### Generation of cell lines expressing fluorescent FLAM8

For EYFP tagging of FLAM8 at the C- and N-terminal ends, p3329 and p2675 plasmids were used, respectively (Kelly et al., 2007). These vectors carry different *FLAM8* gene fragments, corresponding to *FLAM8* ORF nucleotides 8892-9391 and 4-489 in the p3329 and p2675 plasmids, respectively. Both vectors were linearized with NruI and BamHI, respectively, prior to nucleofection following a standard protocol (Burkard et al., 2007). Transformants were selected with appropriate puromycin concentrations (1 or 0.1 μg/ml for PCF or BSF). Clonal populations were obtained by limiting dilution. Cell culture growth was monitored with an automatic Muse cell analyser (Merck Millipore, Paris, France). Neither the FLAM8 targeting to the tip nor the flagellum length were affected by the presence of any EYFP-tags in PCF cells (Fig. S1, A-B).

### Tsetse fly maintenance, infection and dissection

*Glossina morsitans morsitans* tsetse flies were maintained in Roubaud cages at 27°C and 70% hygrometry and fed through a silicone membrane with fresh mechanically defibrinated sheep blood. Teneral males (between 24 h and 72 h post-emergence) were allowed to ingest parasites in culture medium during their first meal through a silicone membrane. Here, we used freshly *in vitro* differentiated stumpy BSF trypanosomes resuspended at 2×10^6^ cells/ml in complete SDM-79 medium supplemented with 10 mM glutathione prior to infection (MacLeod et al., 2007).

Flies were starved for at least 24 hours before being dissected. Different time points were selected for fly dissection: i) 3 days for early isolation of PCF from the posterior midgut (PMG) region; ii) 11 days for recovery of PCF and long trypomastigotes (LT) from the anterior midgut (AMG); iii) 14 days for isolation of all trypomastigote parasites from the midgut and cardia; and, iv) 30 days for isolation of all stages from the MG, PV and salivary glands (SG). For PMG and AMG isolation, the entire midgut was cut into three subsections including the PMG close to the hindgut, the middle part of the MG until the appearance of the bacteriome and the thin AMG that runs from the cardia to the bacteriome. They were isolated accordingly and placed in a drop of phosphate buffer saline (PBS). For recovery of all tsetse organs, after rapid isolation of the salivary glands in a first drop of PBS, whole tsetse alimentary tracts, from the distal part of the foregut to the Malpighian tubules, were dissected and arranged lengthways in another drop of PBS as previously described (Rotureau et al., 2011; Rotureau et al., 2012). Isolated organs were then either scrutinized under a microscope at 40x magnification and imaged, or individually dilacerated, dried on the slide and further treated for immunofluorescence (Rotureau et al., 2014a).

### Mice infection and ethical statements

Seven-week old male BALB/c mice (substrain BALB/cAnNRj) were purchased from Janvier Laboratory (France) and used as models for experimental infection. This study was conducted in strict accordance with the recommendations from the Guide for the Care and Use of Laboratory Animals of the European Union (European Directive 2010/63/UE) and the French Government. The protocol was approved by the “Comité d’éthique en expérimentation animale de l’Institut Pasteur” CETEA 89 (Permit number: 2012-0043 and 2016-0017) and undertaken in compliance with the Institut Pasteur Biosafety Committee (protocol CHSCT 12.131). BR is authorized to perform experiments on vertebrate animals (license #A-75-2035) and is responsible for all the experiments conducted personally or under his supervision. For *in vivo* infections, two BALB/c mice were injected intraperitoneally (IP) with 10^5^ SL BSF parasites, washed in TDB and resuspended in 100 μl PBS prior animal inoculation. Parasitemia was determined daily from tail bleeds. At the first peak of parasitemia, blood containing a mixture of SL and ST parasites was sampled and used to induce the *ex vivo* differentiation of ST parasites into PCF trypanosomes.

### Immunofluorescence analysis (IFA)

Cultured parasites were washed twice in SDM-79 medium without serum or in TDB for PCF and BSF, respectively, and spread directly onto poly-L-lysine coated slides. Both methanol, paraformaldehyde/methanol and cytoskeleton fixation protocols were used for IFA. For methanol fixation, slides were air-dried for 10 min, fixed in methanol at −20°C for 5 min and rehydrated for 20 min in PBS. For PFA/methanol, spread cells were left for 10 min to settle prior to treatment with 1 volume 4% PFA solution in PBS at pH 7. After 5 min, adherent PFA-treated cells were washed briefly in PBS before being fixed with methanol at −20°C for an additional 5 min followed by a rehydration step in PBS for 15 min. For IFA on cytoskeletons, 40 μl of TDB washed cells were spread on a poly-lysine slide. After sedimentation in humid atmosphere, cells were treated for 5 min by addition of 70 μl of PIPES pH 6.9 MgCl_2_, 1 mM 0.4% NP40, then washed twice in PIPES pH 6.9 MgCl_2_ 1 mM and fixed at −20°C in methanol for 5 min. Cells were rehydrated in PBS for 15 min prior to immunodetection by IFA.

For immunodetection, slides were incubated for 1 h at 37°C with the appropriate dilution of the first antibody in 0.1% BSA in PBS. After 3 consecutive 5 min washes in PBS, species and subclass-specific secondary antibodies coupled to the appropriate fluorochrome (Alexa 488, Cy3, Cy5 from Jackson ImmunoResearch, UK) were diluted 1/400 in PBS containing 0.1% BSA and were applied for 1 h at 37°C. After washing in PBS as indicated above, slides were finally stained with 4’,6-diamidino-2-phenylindole (DAPI, 1 μg/ml) for visualization of kinetoplast and nuclear DNA content, and mounted under cover slips with ProLong antifade reagent (Invitrogen), as previously described (Rotureau et al., 2011).

Slides were observed under an epifluorescence DMI4000 microscope (Leica, Germany) with a 100x objective (NA 1.4), an EL6000 (Leica, Germany) as light excitation source and controlled by the Micro-Manager V1.4.22 software (NIH) and images were acquired using a Hamamatsu ORCA-03G or Prime 95B (Photometrics, USA) cameras. Images were analysed with ImageJ V1.8.0 (NIH).

Structured Illumination Microscopy was performed on a Zeiss LSM 780 Elyra PS1 microscope (Carl Zeiss, Germany) using Plan-Apochromat 63Å~/1.4 oil objective with a 1.518 refractive index oil (Carl Zeiss, Germany). The fluorescence signal was detected on an EMCCD Andor Ixon 887 1 K. SIM images were processed with the ZEN software and corrected for chromatic aberration using 100-nm TetraSpeck microspheres (ThermoFisher Scientific). The SIMcheck plugin in imageJ was used to analyze the quality of the acquisition and the processing in order to optimize parameters for resolution, signal-to-noise ratio, and reconstruction pattern.

Primary antibodies were: mAb25 (anti-mouse IgG2a, 1:10), a monoclonal antibody used as a flagellum marker as it specifically recognizes the axoneme protein *Tb*SAXO1 (Dacheux et al., 2012); L8C4 (anti-mouse IgG1, 1:10) (Kohl et al., 1999), a monoclonal antibody that recognizes the PFR2 protein and served as a paraflagellar rod marker; an anti-GFP rabbit serum (Invitrogen #A6455, 1:300) to label the EYFP-tagged versions of FLAM8; a rabbit anti-FTZC (1:1000) which recognizes the Flagellar Transition Zone Component protein in the transzition zone (Bringaud et al., 2000); anti-AK antiserum against *T. cruzi* arginine kinase (anti-mouse IgG, 1:500) (Subota et al., 2014). FLAM8 was detected using anti-FLAM8 rabbit serum (1:500). Specifically, to generate the polyclonal anti-FLAM8 antibody, rabbits were immunised with His-tagged recombinant protein (encoding a 151 amino acid sequence repeated twice in the FLAM8 protein; corresponding to amino acids 442-592 and 593-743). Recombinant protein was expressed in *E. coli* and purified to homogeneity using Immobilised Metal Affinity Chromatography (IMAC) and Anion Exchange Chromatography. Stumpy BSF and early PCF stages were identified at the molecular level with a rabbit polyclonal anti-PAD1 antibody (kindly provided by Keith Matthews, University of Edinburgh; dilution 1:300) and an anti-EP procyclin (Cedarlane^®^, Canada; clone TBRP1/247, antimouse IgG1, 1:500), respectively. When required, sequential incubation with antibodies were performed by means of a saturation step containing 1 % BSA in PBS.

### Measurements, normalization and statistical analyses

Normalization of fluorescent signals was carried out by the parallel adjustments of the raw integrated density and/or minimum signal and/or maximum signal in all the images to be compared in ImageJ V1.8.0 (NIH). For flagellum length measurements, the mAb25 staining of the axoneme was taken as reference using ImageJ. For clarity purposes, the brightness and contrast of several pictures were adjusted after their analysis in accordance with editorial policies. Statistical analyses and plots were performed with XLSTAT 2019.2.01 (Addinsoft) in Excel 2016 (Microsoft) or Prism V8.2.1 (GraphPad). Statistical analyses include linear regression analyses for fluorescence intensity vs. parasite density at 95% confidence. The number of samples analyzed for each experiment is indicated in figure legends.

## Acknowledgements

We thank M. Bonhivers, L. Kohl, D. Robinson, K. Matthews and K. Gull for providing various plasmids and antibodies. We are thankful to the IP BioImaging Plateforme for providing access to their equipment. We acknowledge the generous supply of tsetse pupae from the International Atomic Energy Agency, Vienna. We thank Christelle Travaillé and Aline Crouzols for their help in fly infections. We also thank M. Boshart for his critical reading of the manuscript.

## Author contributions

ECA, SB, PB and BR designed the study. ECA, SB, FEB and AS performed the experiments. ECA, SB, PGMK, PB and BR analysed the data. ECA and BR wrote the manuscript. SB, PGMK and PB reviewed the manuscript.

## References

Abeywickrema, M., Vachova, H., Farr, H., Mohr, T., Wheeler, R. J., Lai, D. H., Vaughan, S., Gull, K., Sunter, J. D. and Varga, V. (2019). Non-equivalence in old-and new-flagellum daughter cells of a proliferative division in *Trypanosoma brucei*. Mol Microbiol 112, 1024–1040.

Acosta-Serrano, A., Vassella, E., Liniger, M., Kunz Renggli, C., Brun, R., Roditi, I. and Englund, P. T. (2001). The surface coat of procyclic *Trypanosoma brucei:* programmed expression and proteolytic cleavage of procyclin in the tsetse fly. Proc Natl Acad Sci U S A 98, 1513–8.

Barquilla, A., Saldivia, M., Diaz, R., Bart, J. M., Vidal, I., Calvo, E., Hall, M. N. and Navarro, M. (2012). Third target of rapamycin complex negatively regulates development of quiescence in *Trypanosoma brucei*. Proc Natl Acad Sci U S A 109, 14399–404.

Bastin, P., MacRae, T. H., Francis, S. B., Matthews, K. R. and Gull, K. (1999). Flagellar morphogenesis: protein targeting and assembly in the paraflagellar rod of trypanosomes. Mol Cell Biol 19, 8191–200.

Bertiaux, E., Morga, B., Blisnick, T., Rotureau, B. and Bastin, P. (2018). A Grow-and-Lock Model for the Control of Flagellum Length in Trypanosomes. Curr Biol 28, 3802–3814 e3.

Bloch, M. A. and Johnson, K. A. (1995). Identification of a molecular chaperone in the eukaryotic flagellum and its localization to the site of microtubule assembly. Journal of cell science 108(Pt 11), 3541–5.

Bringaud, F., Robinson, D. R., Barradeau, S., Biteau, N., Baltz, D. and Baltz, T. (2000). Characterization and disruption of a new *Trypanosoma brucei* repetitive flagellum protein, using double-stranded RNA inhibition. Mol Biochem Parasitol 111, 283–97.

Broadhead, R., Dawe, H. R., Farr, H., Griffiths, S., Hart, S. R., Portman, N., Shaw, M. K., Ginger, M. L., Gaskell, S. J., McKean, P. G. et al. (2006). Flagellar motility is required for the viability of the bloodstream trypanosome. Nature 440, 224–7.

Brun, R. and Schonenberger. (1979). Cultivation and *in vitro* cloning or procyclic culture forms of *Trypanosoma brucei* in a semi-defined medium. Short communication. Acta Trop 36, 289–92.

Buisson, J., Chenouard, N., Lagache, T., Blisnick, T., Olivo-Marin, J. C. and Bastin, P. (2013). Intraflagellar transport proteins cycle between the flagellum and its base. J Cell Sci 126, 327–38.

Chan, K. Y. and Ersfeld, K. (2010). The role of the Kinesin-13 family protein TbKif13-2 in flagellar length control of *Trypanosoma brucei*. Mol Biochem Parasitol 174, 137–40.

Czichos, J., Nonnengaesser, C. and Overath, P. (1986). Trypanosoma brucei: cis-aconitate and temperature reduction as triggers of synchronous transformation of bloodstream to procyclic trypomastigotes in vitro. Experimental parasitology 62, 283–291.

Dacheux, D., Landrein, N., Thonnus, M., Gilbert, G., Sahin, A., Wodrich, H., Robinson, D. R. and Bonhivers, M. (2012). A MAP6-related protein is present in protozoa and is involved in flagellum motility. PLoS One 7, e31344.

Dean, S., Marchetti, R., Kirk, K. and Matthews, K. R. (2009). A surface transporter family conveys the trypanosome differentiation signal. Nature 459, 213–7.

Engstler, M. and Boshart, M. (2004). Cold shock and regulation of surface protein trafficking convey sensitization to inducers of stage differentiation in *Trypanosoma brucei*. Genes Dev 18, 2798–811.

Field, M. C. and Carrington, M. (2009). The trypanosome flagellar pocket. Nat Rev Microbiol 7, 775–86.

Fort, C., Bonnefoy, S., Kohl, L. and Bastin, P. (2016). Intraflagellar transport is required for the maintenance of the trypanosome flagellum composition but not its length. J Cell Sci 129, 3026–41.

Gibson, W. and Bailey, M. (2003). The development of *Trypanosoma brucei* within the tsetse fly midgut observed using green fluorescent trypanosomes. Kinetoplastid Biol Dis 2, 1.

Hirumi, H. and Hirumi, K. (1989). Continuous cultivation of *Trypanosoma brucei* blood stream forms in a medium containing a low concentration of serum protein without feeder cell layers. J Parasitol 75, 985–9.

Hoog, J. L., Lacomble, S., O’Toole, E. T., Hoenger, A., McIntosh, J. R. and Gull, K. (2014). Modes of flagellar assembly in *Chlamydomonas reinhardtii* and Trypanosoma brucei. Elife 3, e01479.

Hughes, L., Towers, K., Starborg, T., Gull, K. and Vaughan, S. (2013). A cellbody groove housing the new flagellum tip suggests an adaptation of cellular morphogenesis for parasitism in the bloodstream form of *Trypanosoma brucei*. J Cell Sci 126, 5748–57.

Kelly, S., Reed, J., Kramer, S., Ellis, L., Webb, H., Sunter, J., Salje, J., Marinsek, N., Gull, K., Wickstead, B. et al. (2007). Functional genomics in *Trypanosoma brucei:* a collection of vectors for the expression of tagged proteins from endogenous and ectopic gene loci. Mol Biochem Parasitol 154, 103–9.

Kohl, L., Robinson, D. and Bastin, P. (2003). Novel roles for the flagellum in cell morphogenesis and cytokinesis of trypanosomes. EMBO J 22, 5336–46.

Kohl, L., Sherwin, T. and Gull, K. (1999). Assembly of the paraflagellar rod and the flagellum attachment zone complex during the *Trypanosoma brucei* cell cycle. J Eukaryot Microbiol 46, 105–9.

Le Ray, D., Barry, J. D., Easton, C. and Vickerman, K. (1977). First tsetse fly transmission of the “AnTat” serodeme of *Trypanosoma brucei*. Ann Soc Belg Med Trop 57, 369–81.

Lemos, M., Mallet, A., Bertiaux, E., Imbert, A., Rotureau, B. and Bastin, P. (2020). Timing and original features of flagellum assembly in trypanosomes during development in the tsetse fly. Parasit Vectors 13, 169.

Liu, W., Apagyi, K., McLeavy, L. and Ersfeld, K. (2010). Expression and cellular localisation of calpain-like proteins in *Trypanosoma brucei*. Molecular and biochemical parasitology 169, 20–6.

MacGregor, P., Szoor, B., Savill, N. J. and Matthews, K. R. (2012). Trypanosomal immune evasion, chronicity and transmission: an elegant balancing act. Nat Rev Microbiol 10, 431–8.

MacLeod, E. T., Maudlin, I., Darby, A. C. and Welburn, S. C. (2007). Antioxidants promote establishment of trypanosome infections in tsetse. Parasitology 134, 827–831.

Oberholzer, M., Langousis, G., Nguyen, H. T., Saada, E. A., Shimogawa, M. M., Jonsson, Z. O., Nguyen, S. M., Wohlschlegel, J. A. and Hill, K. L. (2011). Independent analysis of the flagellum surface and matrix proteomes provides insight into flagellum signaling in mammalian-infectious *Trypanosoma brucei*. Mol Cell Proteomics 10, M111 010538.

Ooi, C. P. and Bastin, P. (2013). More than meets the eye: understanding *Trypanosoma brucei* morphology in the tsetse. Front Cell Infect Microbiol 3, 71.

Ooi, C. P., Rotureau, B., Gribaldo, S., Georgikou, C., Julkowska, D., Blisnick, T., Perrot, S., Subota, I. and Bastin, P. (2015). The Flagellar Arginine Kinase in *Trypanosoma brucei* Is Important for Infection in Tsetse Flies. PLoS One 10, e0133676.

Pedersen, L. B., Geimer, S. and Rosenbaum, J. L. (2006). Dissecting the molecular mechanisms of intraflagellar transport in *Chlamydomonas*. Curr Biol 16, 450–9.

Pereira, C. A., Alonso, G. D., Torres, H. N. and Flawia, M. M. (2002). Arginine kinase: a common feature for management of energy reserves in African and American flagellated trypanosomatids. J Eukaryot Microbiol 49, 82–5.

Reiter, J. F., Blacque, O. E. and Leroux, M. R. (2012). The base of the cilium: roles for transition fibres and the transition zone in ciliary formation, maintenance and compartmentalization. EMBO Rep 13, 608–18.

Robinson, D. R. and Gull, K. (1991). Basal body movements as a mechanism for mitochondrial genome segregation in the trypanosome cell cycle. Nature 352, 731–3.

Roditi, I., Carrington, M. and Turner, M. (1987). Expression of a polypeptide containing a dipeptide repeat is confined to the insect stage of *Trypanosoma brucei*. Nature 325, 272–4.

Roditi, I., Schumann, G. and Naguleswaran, A. (2016). Environmental sensing by African trypanosomes. Curr Opin Microbiol 32, 26–30.

Rotureau, B., Blisnick, T., Subota, I., Julkowska, D., Cayet, N., Perrot, S. and Bastin, P. (2014a). Flagellar adhesion in *Trypanosoma brucei* relies on interactions between different skeletal structures in the flagellum and cell body. J Cell Sci 127, 204–15.

Rotureau, B., Morales, M. A., Bastin, P. and Spath, G. F. (2009). The flagellum-MAP kinase connection in Trypanosomatids: a key sensory role in parasite signaling and development? Cell Microbiol 11, 710–18.

Rotureau, B., Ooi, C. P., Huet, D., Perrot, S. and Bastin, P. (2014b). Forward motility is essential for trypanosome infection in the tsetse fly. Cell Microbiol 16, 425–33.

Rotureau, B., Subota, I. and Bastin, P. (2011). Molecular bases of cytoskeleton plasticity during the *Trypanosoma brucei* parasite cycle. Cell Microbiol 13, 705–16.

Rotureau, B., Subota, I., Buisson, J. and Bastin, P. (2012). A new asymmetric division contributes to the continuous production of infective trypanosomes in the tsetse fly. Development 139, 1842–50.

Rotureau, B. and Van Den Abbeele, J. (2013). Through the dark continent: African trypanosome development in the tsetse fly. Front Cell Infect Microbiol 3, 53.

Saada, E. A., Kabututu, Z. P., Lopez, M., Shimogawa, M. M., Langousis, G., Oberholzer, M., Riestra, A., Jonsson, Z., Wohlschlegel, J. A. and Hill, K. L. (2014). Insect stage-specific receptor adenylate cyclases are localized to distinct subdomains of the *Trypanosoma brucei* flagellar membrane. Eukaryot Cell.

Sharma, R., Peacock, L., Gluenz, E., Gull, K., Gibson, W. and Carrington, M. (2008). Asymmetric cell division as a route to reduction in cell length and change in cell morphology in trypanosomes. Protist 159, 137–51.

Sherwin, T. and Gull, K. (1989). The cell division cycle of *Trypanosoma brucei brucei:* timing of event markers and cytoskeletal modulations. Philos Trans R Soc Lond B Biol Sci 323, 573–88.

Sherwin, T., Schneider, A., Sasse, R., Seebeck, T. and Gull, K. (1987). Distinct localization and cell cycle dependence of COOH terminally tyrosinolated alpha-tubulin in the microtubules of *Trypanosoma brucei brucei*. J Cell Biol 104, 439–46.

Shimogawa, M. M., Ray, S. S., Kisalu, N., Zhang, Y., Geng, Q., Ozcan, A. and Hill, K. L. (2018). Parasite motility is critical for virulence of African trypanosomes. Sci Rep 8, 9122.

Smith, T. K., Bringaud, F., Nolan, D. P. and Figueiredo, L. M. (2017). Metabolic reprogramming during the *Trypanosoma brucei* life cycle. F1000Res 6.

Subota, I., Julkowska, D., Vincensini, L., Reeg, N., Buisson, J., Blisnick, T., Huet, D., Perrot, S., Santi-Rocca, J., Duchateau, M. et al. (2014). Proteomic analysis of intact flagella of procyclic *Trypanosoma brucei* cells identifies novel flagellar proteins with unique sub-localisation and dynamics. Mol Cell Proteomics.

Sunter, J. D., Benz, C., Andre, J., Whipple, S., McKean, P. G., Gull, K., Ginger, M. L. and Lukes, J. (2015). Modulation of flagellum attachment zone protein FLAM3 and regulation of the cell shape in *Trypanosoma brucei* life cycle transitions. J Cell Sci 128, 3117–30.

Szoor, B., Ruberto, I., Burchmore, R. and Matthews, K. R. (2010). A novel phosphatase cascade regulates differentiation in *Trypanosoma brucei* via a glycosomal signaling pathway. Genes Dev 24, 1306–16.

Taylor, A. E. and Godfrey, D. G. (1969). A new organelle of bloodstream salivarian trypanosomes. J Protozool 16, 466–70.

Tetley, L. and Vickerman, K. (1985). Differentiation in *Trypanosoma brucei*: host-parasite cell junctions and their persistence during acquisition of the variable antigen coat. J Cell Sci 74, 1–19.

Urbaniak, M. D., Martin, D. M. and Ferguson, M. A. (2013). Global quantitative SILAC phosphoproteomics reveals differential phosphorylation is widespread between the procyclic and bloodstream form lifecycle stages of *Trypanosoma brucei*. J Proteome Res 12, 2233–44.

Urwyler, S., Studer, E., Renggli, C. K. and Roditi, I. (2007). A family of stagespecific alanine-rich proteins on the surface of epimastigote forms of *Trypanosoma brucei*. Mol Microbiol 63, 218–28.

Van Den Abbeele, J., Claes, Y., van Bockstaele, D., Le Ray, D. and Coosemans, M. (1999). Trypanosoma brucei spp. development in the tsetse fly: characterization of the post-mesocyclic stages in the foregut and proboscis. Parasitology 118, 469–78.

Varga, V., Moreira-Leite, F., Portman, N. and Gull, K. (2017). Protein diversity in discrete structures at the distal tip of the trypanosome flagellum. Proceedings of the National Academy of Sciences of the United States of America 114, E6546–E6555.

Woodward, R., Carden, M. J. and Gull, K. (1995). Immunological characterization of cytoskeletal proteins associated with the basal body, axoneme and flagellum attachment zone of *Trypanosoma brucei*. Parasitology 111(Pt 1), 77–85.

Woodward, R. and Gull, K. (1990). Timing of nuclear and kinetoplast DNA replication and early morphological events in the cell cycle of *Trypanosoma brucei*. J Cell Sci 95(Pt 1), 49–57.

Woolley, D., Gadelha, C. and Gull, K. (2006). Evidence for a sliding-resistance at the tip of the trypanosome flagellum. Cell Motil Cytoskeleton 63, 741–6.

